## Supplemental Figures for "Dynamin 1xA interacts with Endophilin A1 via its spliced long C-terminus for ultrafast endocytosis"

**Figure S1**

A

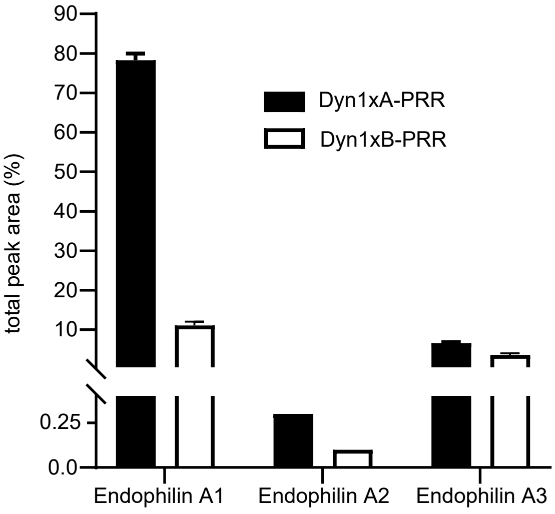


B

Endophilin A1

1. K.VGGAEGTKLDDDFKEMER.K [21, 38]
2. R.AVMEIMTK.T [46, 53]
3. K.LSMINTMSK.I [68, 76]
4. K.QNFIDPLQNLHDK.D [137, 149]
5. K.QAVQILQQVTVR.L [228, 239]
6. R.ALYDFEPENEGELGFK.E [297, 312]

Endophilin A2

1. K.VGGAEGTKLDDDFREMEK.K [21, 38]
2. K.QNFIDPLQNLCDK.D [137, 149]
3. R.QAVQILEELADK.L [228, 239]
4. K.ITASSSFR.S [283, 290]
5. K.ALYDFEPENDGELGFR.E [313, 328]

Endophilin A3

1. K.ASQLFSEK.I [13, 20]
2. K.ATEYLQPNPAYR.A [54, 65]
3. K.DSLDINVK.Q [129, 136]
4. R.QSTEILQELQNK.L [228, 239]
5. R.IALASQVPR.R [244, 252]
6. R.GLYDFEPENEGELGFK.E [292, 307]

**Figure S1. Mass spectrometry evidence for identification of Endophilin A1 bound to Dyn1xA-PRR**

(A) Rat brain synaptosome lysates were incubated with GST-Dyn1-PRR (either xA or xB) coupled to GSH-sepharose beads. Bound proteins were separated by SDS-PAGE and stained with Coomassie blue. The protein band at 40 kDa from both GST-Dyn1-PRR (either xA or xB) were individually cut and digested using Trypsin/LysC and analysed using a targeted LC-MS/MS SRM analyses. The total amount of each Endophilin protein bound to each Dyn1-PRR was calculated by summing the total peak area for each Q1/Q3 transition to provide the total peak area. The percentage of the total peak area for each protein was then calculated. Endophilin A1 was the predominant protein with a level of Endophilin A3 at 11-fold lower levels, while A2 levels were more than 250-fold lower than A1.

(B) List of unique Endophilin isoform-specific peptides used for SRM assay (rat sequences: Endophilin-A1_sp|O35179|SH3G2, Endophilin-A2-sp|O35964|SH3G1, Endophilin-A3-sp|O35180|SH3G3).

**Figure S2**

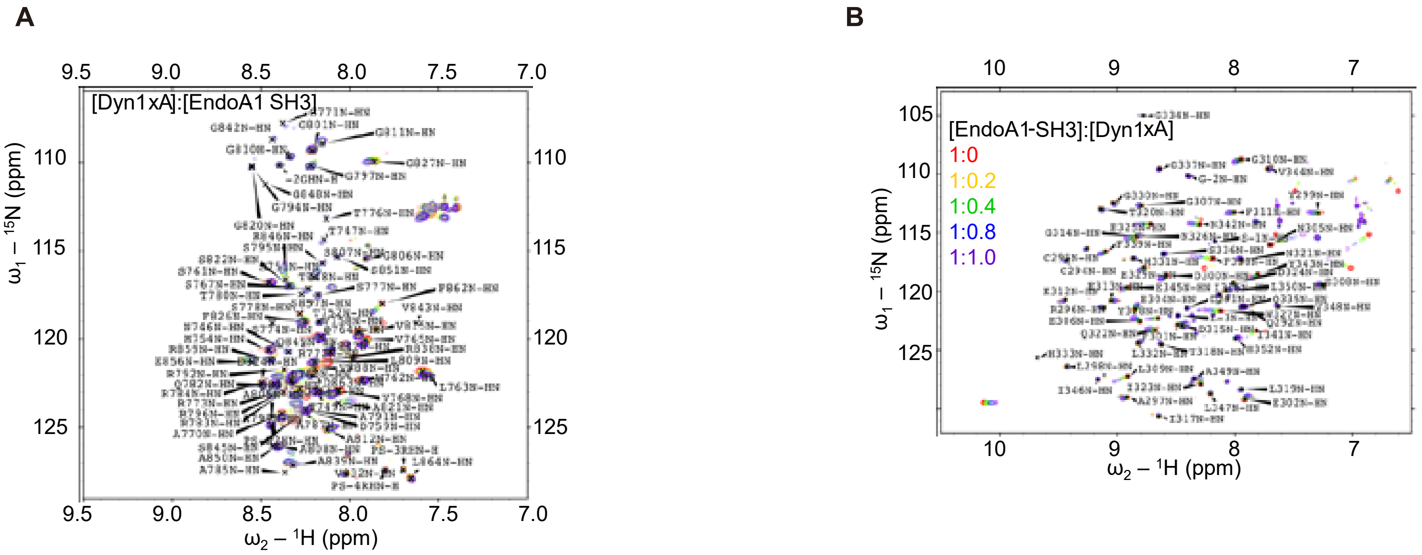
A
[Dyn1xA]:[EndoA1 SH3]
ω1 – 15N (ppm)
ω2 – 1H (ppm)
9.5
9.0
8.5
8.0
7.5
7.0
110
115
120
125
B
[EndoA1-SH3]:[Dyn1xA]
1:0
1:0.2
1:0.4
1:0.8
1:1.0
ω1 – 15N (ppm)
ω2 – 1H (ppm)
10
9
8
7
105
110
115
120
125

**Figure S2. Dynamin 1xA has two distinct binding sites for Endophilin A1 SH3**

(A) Overlaid $^{1}H$-$^{15}N$ HSQC spectra of the isolated [U-$^{15}N$]GST-tagged cleaved Dyn1xA 746-798 recorded upon the titration of the nonlabeled GST-tagged cleaved endophilin SH3 domain.

(B) Overlaid $^{1}H$-$^{15}N$ HSQC spectra of the isolated [U-$^{15}N$] GST-tag cleaved Endophilin A1 SH3 domain recorded upon the titration of the nonlabeled GST-tag cleaved Dyn1xA 746-798.

Figure S3

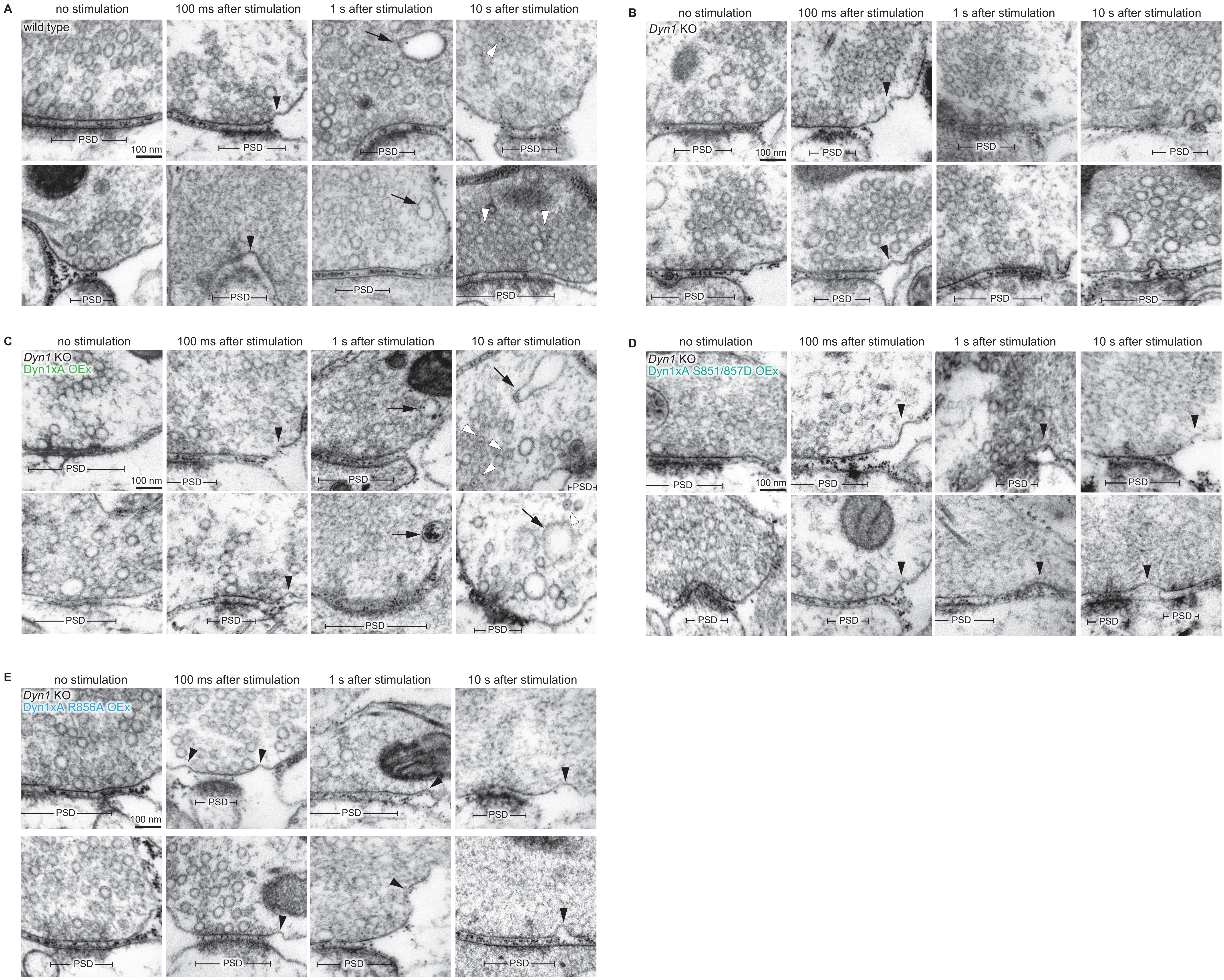
A
no stimulation
100 ms after stimulation
1 s after stimulation
10 s after stimulation
wild type
PSD
100 nm
B
no stimulation
100 ms after stimulation
1 s after stimulation
10 s after stimulation
Dyn1 KO
PSD
100 nm
C
no stimulation
100 ms after stimulation
1 s after stimulation
10 s after stimulation
Dyn1 KO
Dyn1xA OEx
PSD
100 nm
D
no stimulation
100 ms after stimulation
1 s after stimulation
10 s after stimulation
Dyn1 KO
Dyn1xA S851/857D OEx
PSD
100 nm
E
no stimulation
100 ms after stimulation
1 s after stimulation
10 s after stimulation
Dyn1 KO
Dyn1xA R856A OEx
PSD
100 nm

**Figure S3. Additional EM images for Figure 5.**

Example micrographs showing endocytic pits and ferritin-containing endocytic structures at the indicated time points in wild-type neurons, *Dyn1* KO neurons, and *Dyn1* KO neurons overexpressing Dyn1xA (Dyn1xA OEx), Dyn1xA S851/857D (Dyn1xA S851/857D OEx) and Dyn1xA R846A (Dyn1xA R846A OEx). Scale bar: 100 nm.

Figure S4

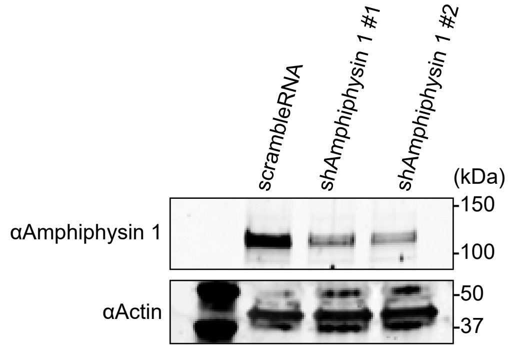
A
scrambleRNA
shAmphiphysin 1 #1
shAmphiphysin 1 #2
(kDa)
αAmphiphysin 1
150
100
αActin
50
37

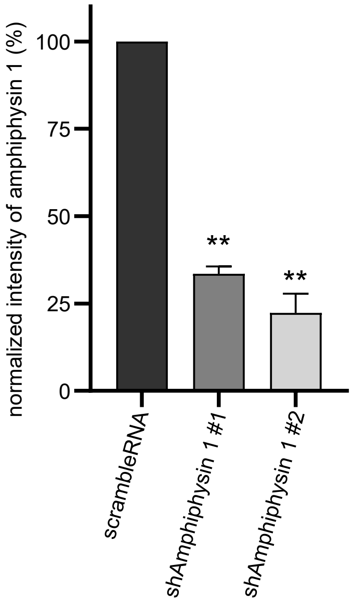
B
normalized intensity of amphiphysin 1 (%)
100
75
50
25
0
**
**
scrambleRNA
shAmphiphysin 1 #1
shAmphiphysin 1 #2

**Figure S4. Evaluation of Amphiphysin 1 knock down.**

(A) An example immunoblotting images showing anti-Amphiphysin 1 or anti-β-Actin antibodies reactions against the lysates of cultured hippocampal neurons infected with lentivirus expressing scramble shRNA and two different sequences of Amphiphysin 1 shRNA.

(B) Normalized signal intensities of Amphiphysin 1 quantified the immunoblotting in (A).

*$p < 0.05$, unpaired t test. The mean and SEM are shown. $n = 2$ independent cultures.

Figure S5

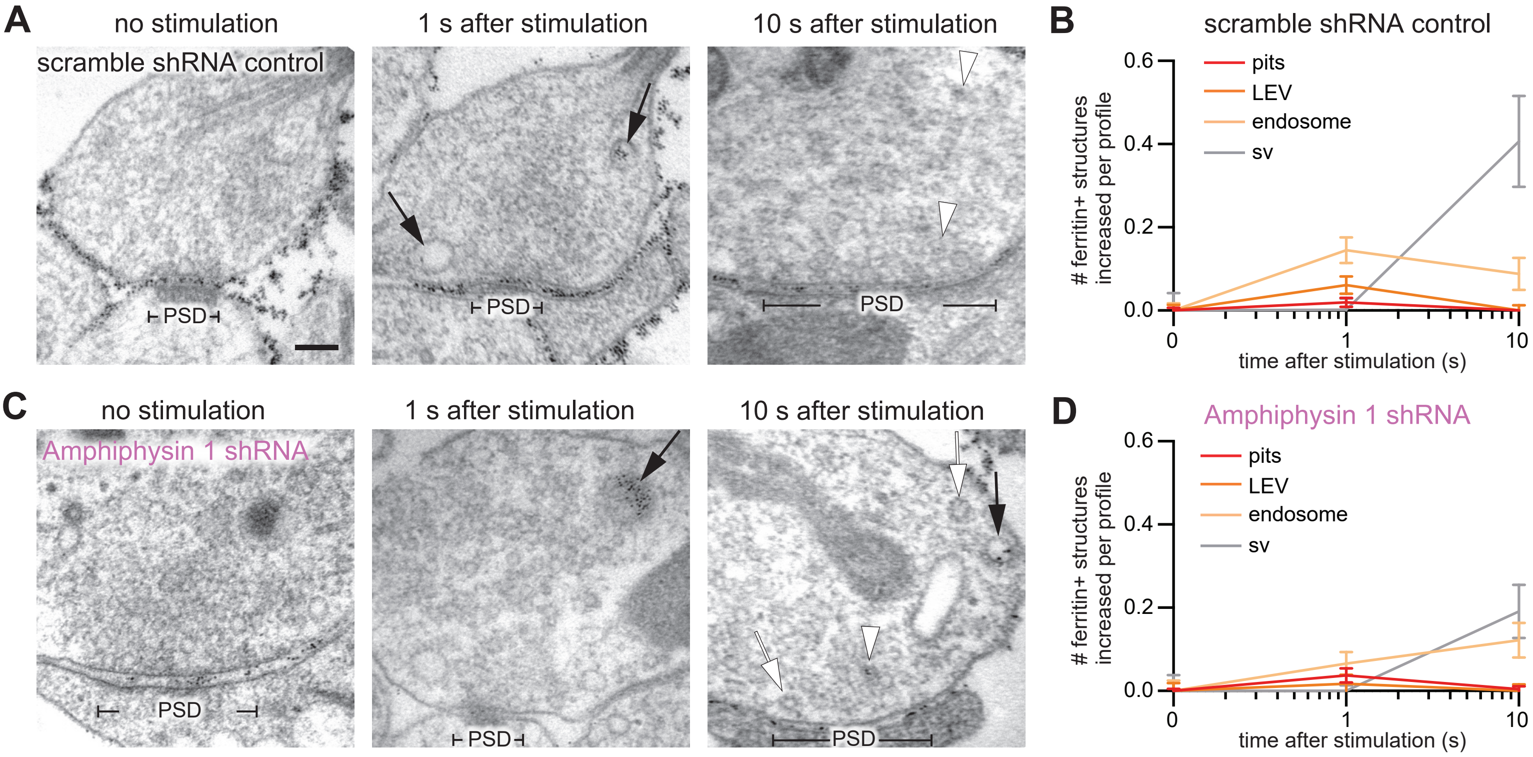

A
no stimulation
scramble shRNA control
PSD
1 s after stimulation
PSD
10 s after stimulation
PSD
B
scramble shRNA control
### ferritin+ structures increased per profile
0.0
0.2
0.4
0.6
pits
LEV
endosome
sv
0
1
10
time after stimulation (s)
C
no stimulation
Amphiphysin 1 shRNA
PSD
1 s after stimulation
PSD
10 s after stimulation
PSD
D
Amphiphysin 1 shRNA
### ferritin+ structures increased per profile
0.0
0.2
0.4
0.6
pits
LEV
endosome
sv
0
1
10
time after stimulation (s)

**Figure S5. Amphiphysin 1 is not essential for ultrafast endocytosis.**

(A and C) Example micrographs showing endocytic pits and ferritin-containing endocytic structures at the indicated time points in neurons expressing scramble RNA (A) and Amphiphysin 1 shRNA (C). Black arrows, ferritin-positive large endocytic vesicles (LEVs) or endosomes; white arrowheads, ferritin-positive synaptic vesicles. Scale bar: 100 nm. PSD, post-synaptic density.

(B and D) Plots showing the increase in the number of each endocytic structure per synaptic profile after a single stimulus in neurons expressing scramble RNA (B) and Amphiphysin 1 shRNA (D). The mean and SEM are shown in each graph.

All data are from two independent experiments from N = 2 cultures prepared and frozen on different days. n = scramble RNA, 436; Amphiphysin 1 shRNA, 609.

Figure S6

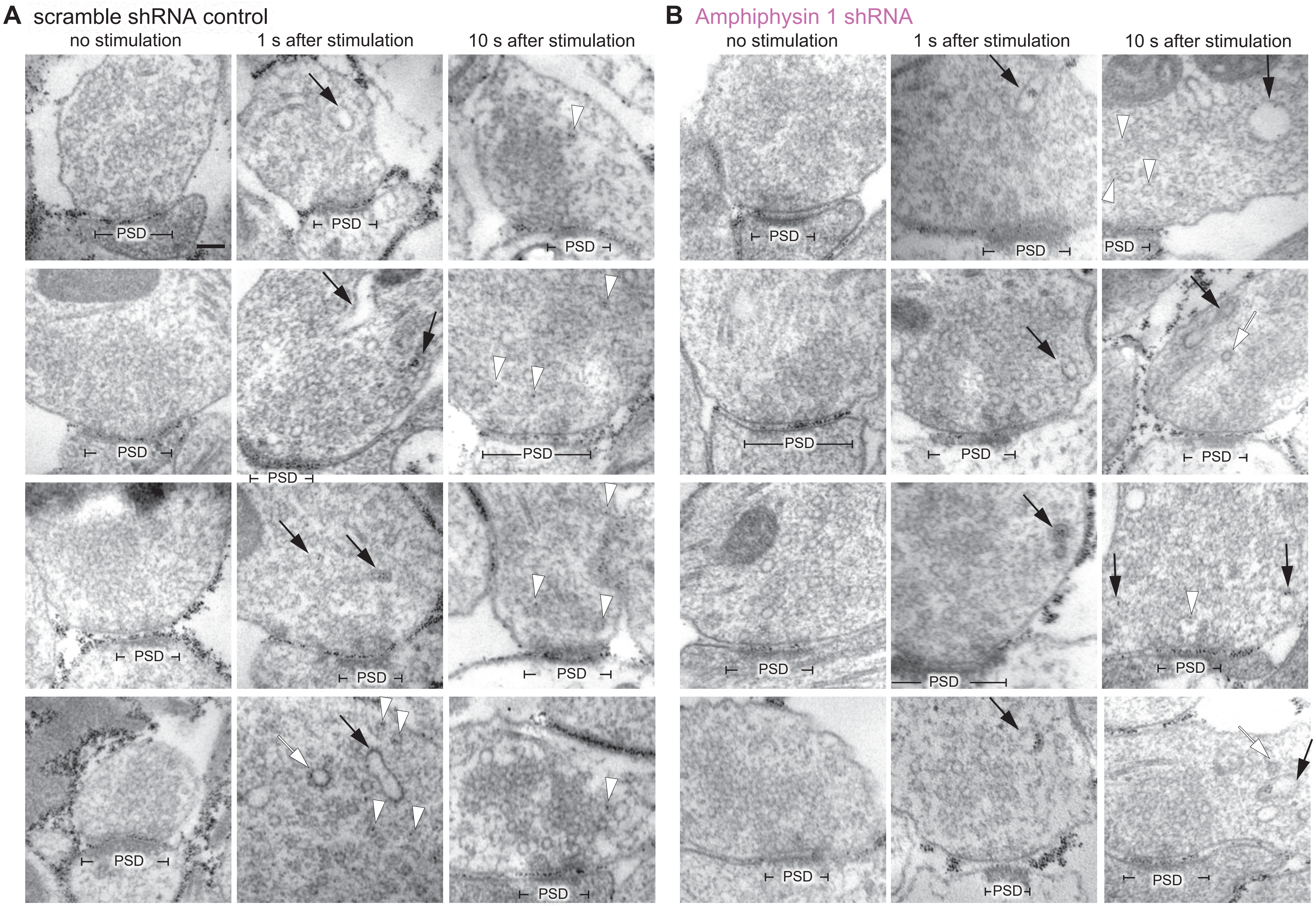
A scramble shRNA control
no stimulation
1 s after stimulation
10 s after stimulation
B Amphiphysin 1 shRNA
no stimulation
1 s after stimulation
10 s after stimulation
PSD

**Figure S6. Additional EM images for Figure S3.**

Example micrographs showing endocytic pits and ferritin-containing endocytic structures at the indicated time points in neurons expressing scramble RNA (A) and Amphiphysin 1 shRNA (C). Black arrows, ferritin-positive large endocytic vesicles (LEVs) or endosomes; white arrow, clathrin-coated vesicle; white arrowheads, ferritin-positive synaptic vesicles. Scale bar: 100 nm. PSD, post-synaptic density.

Figure S7

A

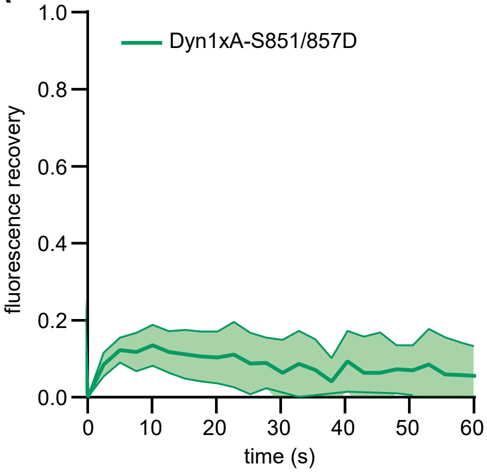

Dyn1xA-S851/857D
fluorescence recovery
1.0
0.8
0.6
0.4
0.2
0.0
0
10
20
30
40
50
60
time (s)

**Figure S7. Dyn1xA-S851/857-GFP puncta are dominated by immobile fraction.**

(A) Examples live images of FRAP experiments of Dyn1xA-S851/857-GFP puncta in presynapse. mCherry-synaptobrevin 2 (mCherry-Syb2) was tandemly expressed to find presynapses. Dyn1xA-S851/857-GFP signals were photobleached at 480 nm. Time indicates after the photobleaching.

(B) Normalized fluorescence recovery of Dyn1xA-S851/857-GFP signals. Fluorescence signals were normalized to just after (0 s) photobleaching. Times indicate after the photobleaching. The median and 95% confidential interval are shown.

n = 18 presynapses from two independent cultures.
